# Resin canal leakage underlying xylem dysfunction in pine wilt disease

**DOI:** 10.1101/2025.06.06.656100

**Authors:** Toshihiro Umebayashi, Kenji Fukuda

## Abstract

During the course of pine wilt disease (PWD), xylem dysfunction is induced (in the absence of water stress) by invasion of pinewood nematodes (PWN) into resin canals, and the infected pine trees wilt. When PWNs are inoculated to Japanese black pine seedlings, massive embolisms develop at the inoculation site. The developmental process of massive embolization to form an embolized zone is not well understood. We studied the development of xylem dysfunction around the inoculation site of one-year-old stems of PWN-infected seedlings using a combination of compact MRI and cryo-SEM. We visualized pathological changes in tracheids and resin canals in the xylem, and found that embolisms occurred in PWN-inoculated seedlings at a negative hydraulic pressure of −0.5 MPa. Resin canals in healthy trees were filled with resin. In PWN-infected seedlings, abnormal embolisms and resin were observed around resin canal near the inoculation wound. Empty tracheids were observed more extensively than resin-filled tracheids. We suggest that abnormal embolisms and resin leakage occur from damages of resin canals. Resin may seep into the tracheids when cell wall of the axial and radial resin canals is crushed. These embolisms under high water potential are unavoidable under natural conditions.

## Introduction

Woody plants lose water through leaf transpiration. Water is resupplied through root uptake and subsequent transport under negative pressure through the vascular conduits (vessels and tracheids) to the leaves. Under water stress, cavitation in conduits arises at the air-water meniscus in the pit aperture (Sperry and Tyree, 1988; Tyree and Zimmermann, 2002), resulting in embolisms within the conduits. The number of embolized conduits increases with increased negative pressure (Sperry and Tyree, 1988), and the relationship between the loss of hydraulic conductivity and the increase in negative pressure is described by a vulnerability curve (Sperry and Tyree, 1988; Sperry and Tyree, 1990; Umebayashi et al., 2016a). In healthy plants, a loss of hydraulic conductivity can be recovered by refilling embolized conduits and/or the formation of new conduits. Although significant increases in losses of hydraulic conductivity occur only under severe water stress in healthy plants, some vascular wilt diseases, such as Dutch elm disease, induce the decrease of sap flow that kill host trees without water stress (Urban and Dvořák, 2014). The developmental process of embolisms in plants affected by wilt diseases is not fully understood.

Pine trees infected with pine wilt disease (PWD) have high mortality rates. PWD induces abrupt wilt death in susceptible pine species, such as Japanese black and red pines (*Pinus thunbergii* Parl. and *P. densiflora* Siebold et Zucc., respectively) (Tamura, 1983; Umebayashi et al., 2016b). The pathogen in PWD is an alien invasive nematode from North America, the pinewood nematode (PWN) *Bursaphelenchus xylophilus* (Steiner et Buhler) Nickle (Tokushige and Kiyohara, 1969; Kiyohara and Tokushige, 1971). Explanations of the wilting mechanism of PWD have been reviewed many times (Mamiya, 1983; Kuroda, 1991; Fukuda, 1997; Kuroda, 2008). According to a common view, PWN can migrate through the axial and radial resin canals in a living host (Mamiya 1983, Kusumoto et al., 2014; Son et al., 2010, 2015). Most PWN individuals remain near the inoculation wound (Mamiya, 1975; Ichihara et al., 2000), but a small proportion of the worms invade the cortical and xylem resin canals during the early stage of PWD development (Mamiya, 1985; Kuroda, 2008), inducing patchy embolisms around the xylem resin canals. Mamiya (1985) suggested that the existence of a large PWN inoculum at the site of entry is necessary for the induction of systemic disease symptom development, and excision of inoculated branches significantly delays symptom development. However, patchy embolisms eventually enlarge and fuse through the whole xylem area, leading to tree death (Kuroda, 1991; Fukuda et al., 2007).

Contrasting theories and speculations about disease symptom development exist. Pathological changes in living cells are detected by PWN activity, but details of the developmental processes during early stages have differed among observers. Causative materials supposed to induce xylem embolisms vary markedly among research groups. Abnormal embolism formation has been explained by (i) resin leakage from resin canals (Sasaki et al., 1984), (ii) synthesis of volatile terpenes by epithelial cells and ray parenchyma cells as a defense reaction (Kuroda, 1991, 2008), and (iii) osmiophilic depositions from ray parenchyma (Nobuchi et al., 1984; Hara and Futai, 2001). Whether these explanations are really related to the occurrence of xylem embolisms in PWD remains unclear because previous studies have not provided detailed or quantitative relationships between increases in the embolization of tracheids and the distribution of the proposed causal agent. Artifactual embolisms may have been induced by stem cutting in earlier works. The quality of anatomical observations may have been reduced by alterations in the distribution of leaked materials in the xylem. Spatial analyses based on non-invasive visualization at the cellular level are required to identify the substance that induces xylem embolisms in PWD.

In recent work, we have used a compact magnetic resonance imaging (CMRI) system to non-destructively monitor the development of embolisms in the stems of seedlings (Umebayashi et al., 2011, 2016a; Fukuda et al., 2015). We studied the developmental process of PWD xylem dysfunction. Using multiple cross-sectional images, we showed that massive embolisms around the inoculation wound in the stem of Japanese black pine seedlings enlarged in all directions (i.e., axially, radially, and tangentially) prior to the development of patchy embolisms in all monitoring positions. Patchy embolisms were detected and increased in any positions, although they hardly expand unlike the massive embolism at the inoculation site. Embolisms did not occur along the ray parenchyma, but were found along axial and radial resin canals (Umebayashi et al., 2016b); PWNs were detected in/around the resin canals after the parenchyma cells around the resin canals had been crushed (Umebayashi et al., 2017). Massive embolisms occurred around the inoculation site, and they enlarged rapidly, but this was not the case for patchy embolisms in other regions. These observations demonstrate the importance of nematode activities around the inoculation site, and may explain delayed symptom development following excision of an inoculated branch (Mamiya, 1985). Although these results shed some light on the mechanism of embolism development, a combination of different methods, such as non-invasive monitoring of xylem water and anatomical analysis at the cellular level, is necessary to identify in detail the causal agents of xylem embolism and the developmental process of massive embolisms (formation and enlargement of the embolized zone) around the inoculation site.

In this study, we aimed to explore the developmental process of massive embolisms in PWN-infected seedlings using a combination of CMRI and cryo-scanning microscopy (cryo-SEM). First, we explored the threshold value of negative pressure under which xylem embolisms developed in infected trees. We used a blackout curtain to control negative pressure in PWN-infected seedlings. Subsequently, we monitored the embolized area around the inoculation wound during increases in negative pressure. Second, using cryo-SEM, we explored at the cellular level the relationship between the distribution of embolized areas and the pathological changes in living cells in/around resin canals after PWN infection. Finally, we considered the most appropriate hypothesis explaining the mechanism of the xylem dysfunction during early stage PWD.

## Materials and methods

### Plant materials

We used 12 seedlings of three-year-old Japanese black pine grown at the Kashiwa campus, the University of Tokyo (Kashiwa City, Chiba, Japan) as experimental material. The seedlings were 0.9–1.2 m tall, and the diameters of one-year-old stems ranged from 1.0 to 1.2 cm.

We inoculated six seedlings on August 9, 2011 and four seedlings on July 18, 2012. Two controls were injected with distilled water in an identical manner on August 9, 2011. A virulent isolate (Ka-4) of PWN was reared on *Botrytis cinerea* Pers. growing at 25°C on a potato dextrose agar plate. PWNs were isolated from the culture medium using the Baermann funnel method. Ten seedlings were inoculated by injecting 0.01 mL of water containing a suspension of 5,000 nematodes through an oblique wound (*ca*. 1.0 cm in length) cut with a razor blade in a central position on the stem surface of a one-year-old main shoot; the inoculum was injected into the current-year xylem of the selected stem (see Umebayashi et al., 2016b).

The xylem water potential was measured during the experiment with a pressure bomb (Model 3000; Soilmoisture Equipment Corp., Santa Barbara, CA, USA). During the experiment, all seedlings were kept in a laboratory at room temperature (*ca*. 27°C), a relative humidity of 50–60%, and a light period from 06:00 to 17:00 each day.

### Relationship between water potential and embolism development under different water stress conditions determined by CMRI

Six inoculated seedlings and two controls were used to monitor the development of xylem embolisms with or without PWN inoculation (Supplemental Fig. S1 A–C). Following PWN inoculation on August 9, 2011, four inoculated and two control seedlings were kept illuminated on the floor of a laboratory (LI 1–4 and LC 1 and 2, see Supplemental Fig. S1 A, B); the other two inoculated seedlings (DI 1 and 2) were kept in darkness (to stop leaf transpiration) by enclosing them within a blackout curtain. At midnight (i.e., in darkness), we monitored the water distributions in the stems using CMRI, and measured the water potential; this procedure prevented an increase in negative pressure. We supplied 500 mL of water to each seedling each morning throughout the experiment.

Water distributions were monitored by CMRI at the same positions 5 and 8 days after inoculation. Two seedlings (DI 1 and 2) were removed from behind the blackout curtain 8 days after PWN inoculation (i.e., August 17); we continued to visualize the development of the embolized areas with increases in negative pressure under illumination. After CMRI monitoring, the stems of all samples were cut underwater at ground level, and a 10-cm-long segment of each stem that included the monitoring site was held underwater. Each of these segments was frozen *in situ* in liquid nitrogen for later cellular-level visualization of the disease symptoms in the xylem using cryo-SEM.

Four seedlings (DWI 1–4) were inoculated on July 18, 2012. Each was tightly enclosed in a plastic bag containing a wet paper towel and misted with water to prevent any transpiration or evaporation; they were then transferred to darkness behind the blackout curtain. Fourteen days after PWN inoculation, two seedlings (DWI 1 and 2) were removed from behind the blackout curtain. The water distributions of transverse sections through each inoculation site were regularly monitored for (i) water loss in the xylem and (ii) increases in negative pressure. We regularly measured water potential during CMRI monitoring. Following CMRI monitoring, frozen stem samples of two seedlings were subjected to the same procedure.

The other two seedlings (DWI 3 and 4) were kept behind the blackout curtain. Frozen samples that had not experienced increases in negative pressure were also taken using the same method.

### CMRI monitoring

The water distribution in the xylem of 8 inoculated and two control seedlings were monitored using a 0.3 Tesla CMRI system (MR Technology, Inc., Tsukuba, Japan). We selected two monitoring positions immediately above and below the inoculation wound (+1 cm and −1 cm positions; see Umebayashi et al. 2016b) for determination of the embolized area proportions; this procedure was not applied to the two DWI seedlings (DWI 1 and 2). The water distribution in transverse sections at the inoculation sites (i.e., at the 0 cm position) in two DWI seedlings were regularly monitored to visualize details of embolism development outward from the inoculation wound. Two-dimensional cross-sectional images were captured to detect the xylem water distribution under the same conditions applied in the previous MRI experiment, i.e., T1-weighted spin-echo sequence of repetition time, 500 ms; echo time, 19 ms; slice thickness, 1 mm (see, Umebayashi et al., 2011).

### Analysis of CMRI data

To calculate the area of embolization, MR images were binarized using image processing software (Photoshop 7.0, Adobe Systems, San Jose, CA, USA) at a threshold at which white noise in the background was eliminated. From the number of white pixels before (A_init_) and after inoculation (A), the relative embolized area (REA) was calculated as REA (%) = 100 (1 – A/A_init_).

### Cryo-SEM observations

All frozen samples were stored in liquid nitrogen. To observe histological changes in the xylem, a frozen sample was excised at the position monitored by CMRI, then polished with a sliding microtome in a cold room (–27°C). The block was subsequently transferred to the cold stage of the cryo-SEM (JSM-6390LV, JEOL, Tokyo, Japan). The frozen cut surfaces were etched, and observed uncoated at 3 kV.

## Results

### Relationship between water potential and embolism development

Before inoculation, embolized areas were observed around the latewood tracheids of the previous year’s xylem in all samples (Fig. 1). In both control seedlings, the xylem pressure was ≥–0.4 MPa during the experimental period. New embolisms from the wound at the water injection site were barely visible at both monitoring positions above or below the wound (+1 cm and −1 cm positions).

**Figure 1.**
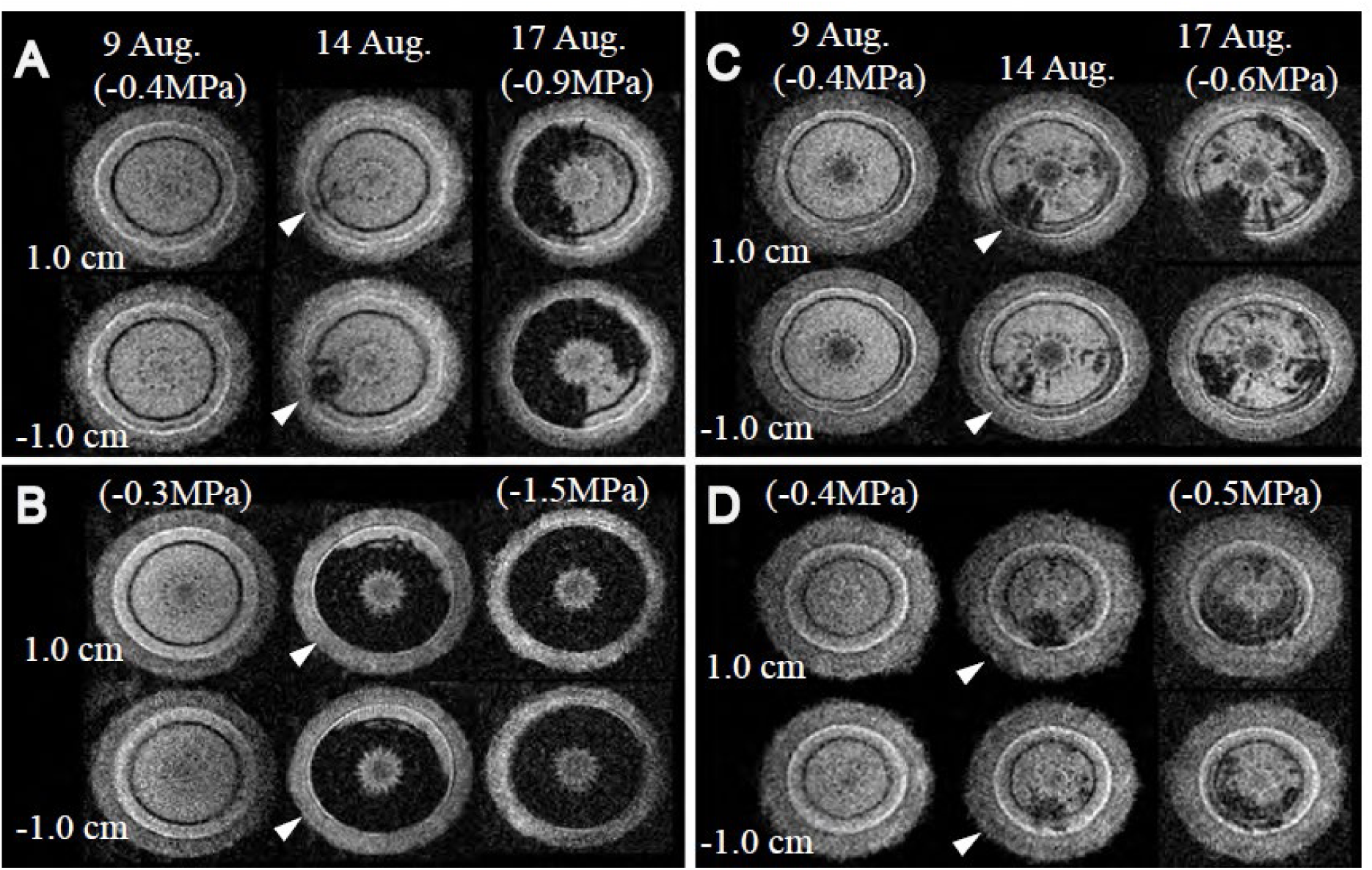
Cross-sectional magnetic resonance images and water potentials obtained at positions 1 cm above and 1 cm below pine wilt nematode (PWN) inoculation wounds on 1-year-old stems of three seedlings held under illumination (A–C) and one seedling kept in darkness (D) from August 9 to August 17. A, LI 1; B, LI 3; C, LI 4; D, DI 1. Arrowheads point to inoculation sites.

In the four PWN-inoculated seedlings exposed to light (LI 1–4), the water potential varied from −0.6 to −1.5 MPa 8 days after PWN inoculation (August 17, 2011; Fig. 1A–C). Among PWN-infected seedlings held in darkness, the water potentials of DI seedlings (DI 1 and 2) were lower (–0.5 MPa, Fig. 1D) than those of DWI seedlings (DWI 1–4) (≥–0.2 MPa). Figure 1 presents examples of the water distribution images obtained by CMRI; the REAs calculated from the images are presented in Figure 2. The REAs of illuminated LI seedlings exceeded those of DI seedlings held in darkness (Fig. 2A). Higher REAs were correlated with low water potentials. In some seedlings, the REA at the −1 cm position exceeded that at the +1 cm position (see Fig. 1A).

**Figure 2.**
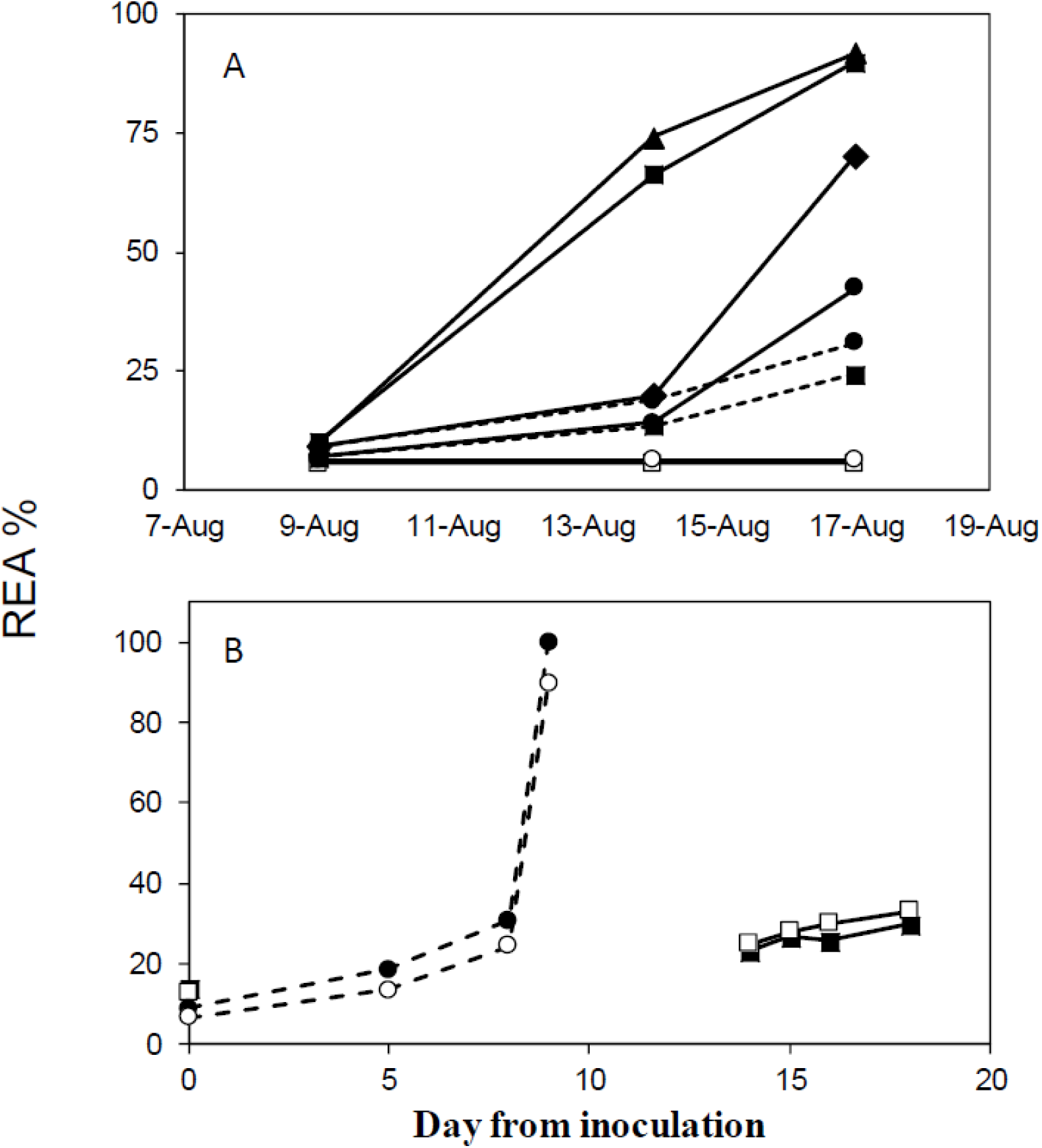
Changes in the relative embolized area (REA) proportions detected by MRI. A, proportions of REA at locations with elevated embolized ratios 1 cm above and 1 cm below the inoculation sites in the period August 9 to August 17, 2012. Open and closed symbols refer to control and PWN-inoculated samples, respectively. Solid lines indicate the four illuminated (LI) seedlings; dashed lines indicate two seedlings held in darkness (DI). Circles, squares, triangles, and diamonds indicate seedling numbers 1, 2, 3, and 4, respectively. B, proportions of REA in seedlings DI 1 and 2 and DWI 1 and 2 from the day of inoculation (MRI observations). Square symbols indicate the two DWI seedlings; circle symbols indicate the two DI seedlings. Open and closed symbols indicate seedling numbers 1 and 2, respectively.

Although high negative pressures were related to the expansive areas of massive embolisms developed at the inoculation wound, massive embolisms were also observed in DI seedlings at the highest water potential (Fig. 1D).

### Spatial analysis of embolized area development by CMRI

The distributions and the shapes of the embolized areas among the four LI seedlings (LI 1–4) were diverse (Fig. 1A–C). Massive embolisms developed in seedlings with large REAs and were observed in both monitoring positions, i.e., lengths greatly exceeded 2 cm (Fig. 1A, B). Patchy embolisms tended to be more abundant in LI seedlings with low REAs and higher water potentials (Fig. 1C). In one illuminated seedling (LI 4), massive embolisms were observed only around the inoculation site; occasional patchy embolisms were also observed at both monitoring positions (Fig. 1C). The areas of patchy embolisms generally developed tangentially from the direction of the inoculation wound 5–8 days after inoculation; the developmental process differed from those of other seedlings (i.e., LI 1–3, Fig. 1A, B). Many small clustered embolisms occurred under high water potential in the seedlings held in darkness (DI 1 and 2) (Fig. 1D). The densely distributed dot-like embolisms in the seedlings held in darkness resembled the area containing both embolized and water-filled tracheids at the inoculation wound, and were different in appearance from both patchy embolisms and massive embolisms in illuminated seedlings. After removal from the blackout curtain (*ca*. 200 min later), the many dot-like embolisms in the two DI seedlings fused together and dramatically expanded tangentially at both monitoring positions (Fig. 3A), and the water potential decreased slightly to −0.7 MPa under illuminated conditions. Finally, the whole area of xylem became embolized at −0.7 MPa *ca*. 15 h (920 min) after removal from the light (Fig. 3A).

**Figure 3.**
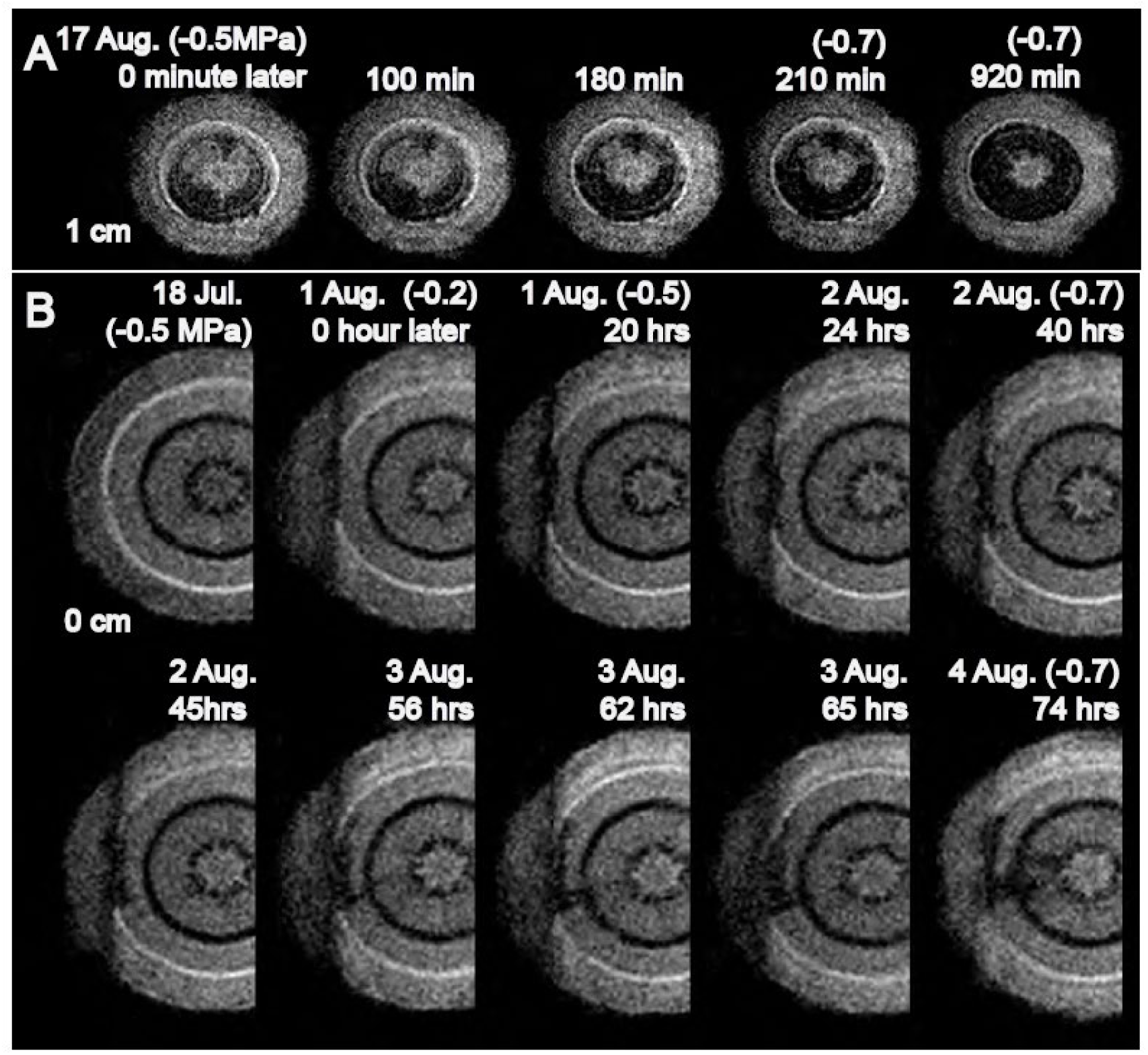
Development of embolized areas in 1-year-old stems of seedlings DI 1 (A) and DWI 1 (B) (i) 1 cm distant from the pine wilt nematode (PWN) inoculation site, and (ii) at the inoculation site before and after transfer to illuminated conditions (data obtained by cross-sectional MRI). Seedling DI 1 (A) was held in darkness in the laboratory behind a blackout curtain (thereby preventing leaf transpiration) through August 17. Seedling DWI 1 (B) was held in darkness in a plastic bag with a wet paper towel and misted with water during the period from July 18 to August 1.

However, two DWI seedlings did present these dot-like embolisms, and the embolized area expanded only slightly in a tangential direction from the inoculation wound at −0.5 MPa *ca*. 20 h after the transfer from darkness to illuminated conditions (Fig. 3B). About 40 h later, the water potential reached −0.7 MPa, and an embolized area developed tangentially from the inoculation wound. About 3 days thereafter, new embolisms were observed in the pith, and the embolized area expanded radially (Fig. 3B, August 4). Although the REA proportions in DI 1 and DI 2 exceeded 80%, the REAs in DWI 1 and 2 approached *ca*. 40% under illumination (Fig. 2B).

### Cellular-level pathological changes in xylem; detection of pinewood nematodes (PWN) in PWN-infected seedlings

In both control seedlings (LC 1 and 2), almost all tracheids were filled with water, except in the latewood of the previous year’s xylem (Fig. 4A). Many resin canals were surrounded by normal epithelial cells and were filled with resin (Fig. 4B, arrowheads). However, most tracheids around the inoculation wound were embolized in the LI seedlings (Figs. 4C, 5A), forming massive embolisms. Within the massive embolisms, epithelial and parenchyma cells around the axial and radial resin canals were often crushed and empty (Figs. 4D, E; 5A). Many resin canals that were partially or fully plugged with expanded epithelial cells (i.e., tylosoid) also occurred in LI seedlings. Tylosoid occurred mostly (i) in patchy embolisms (Fig. 5B), and (ii) in water-filled xylem at the borders of the massive embolisms (Fig. 5C). Whereas most tracheids were empty in the massive embolisms of LI seedlings (Figs. 4C, D, E; 5A), widely distributed resinous substances within the tracheids were observed in the massive embolisms of DI (Fig. 6) and DWI 1 and 2 (Fig. 7). In DWI 1 and 2, many embolized tracheids occurred at the borders of water-filled and resin-filled tracheids (Fig. 7B) distributed around empty resin canals and cavities (Fig. 7A). A mosaic of abundant empty tracheids and resin-filled tracheids was observed in the massive embolisms in DI seedlings (Fig. 6A, B). PWNs were observed in the resin canals in LI and DI seedlings after the epithelial cells had been crushed (Figs. 4E, 5D, 6C), but were rarely observed in the resin-filled resin canals or those occluded by tylosoid.

**Figure 4.**
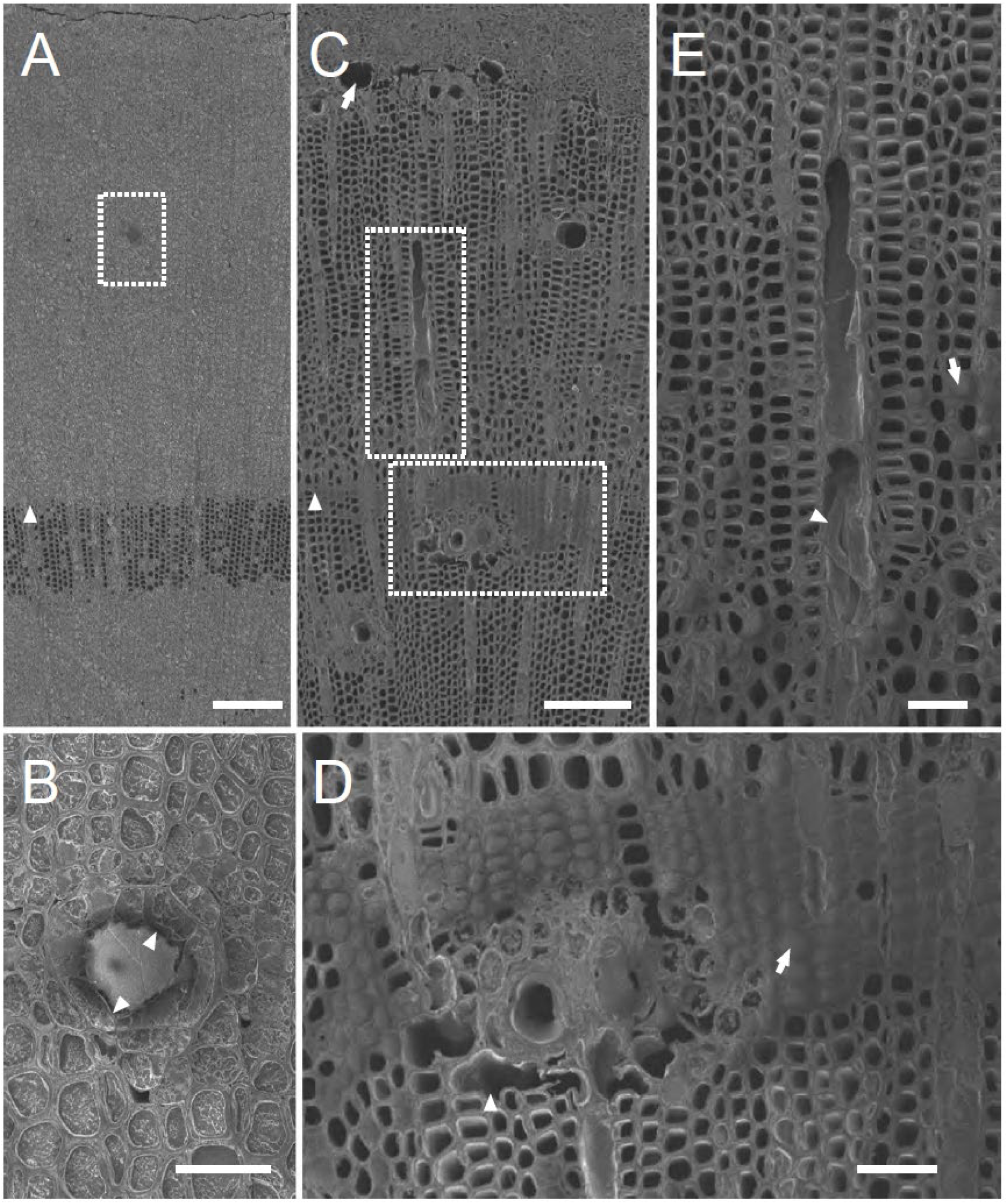
Cryo-scanning microscopy (cryo-SEM) images of cross sections of 1-year-old stems taken 1 cm above the inoculation wound sites in seedlings LC 1 (A, B) and LI 3 (C, D, E). A, water distribution in a cross section of seedling LC 1. The arrowhead points to the annual ring border. The dashed rectangle encloses the extended area shown in B. B, magnified view of an axial resin canal in the current-year xylem of seedling LC 1. Arrowheads point to sound epithelial cells. C, massive embolisms in seedling LI 3. The arrowhead points to the annual ring border. The dashed rectangles enclose the extended areas in D and E. D, magnified view of an axial resin canal in the one-year-old xylem. The arrowhead points to crushed axial parenchyma around the resin canal. The arrow points to resinous substance in the tracheids. E, magnified view of a radial resin canal in the one-year-old xylem. The arrow points to the resinous substance in a tracheid. The arrowhead points to a nematode. Scale bars: A and C, 200 μm; B, D, and E, 50 μm.

**Figure 5.**
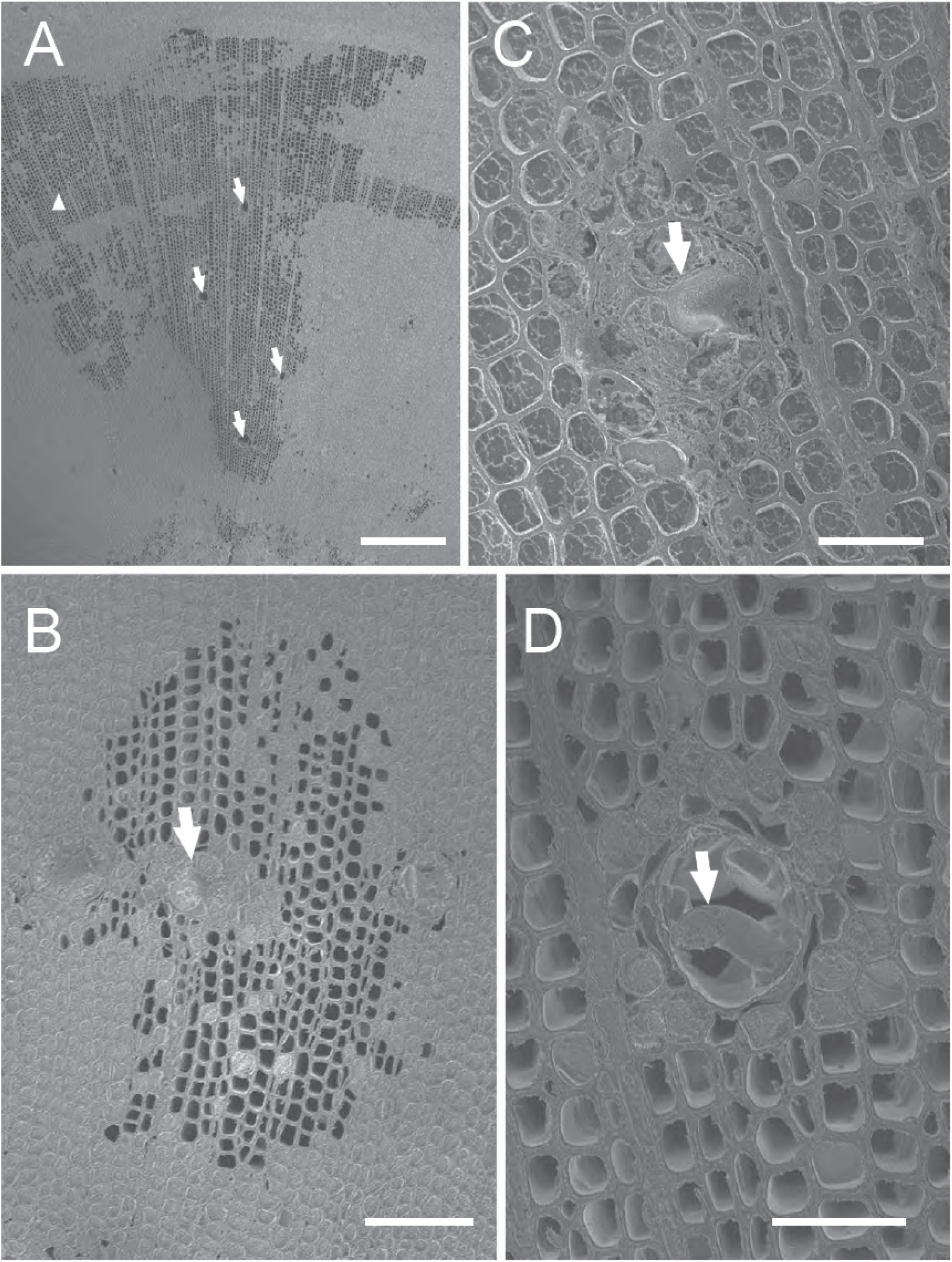
Cryo-SEM images of cross sections of 1-year-old stems 1 cm above the injection sites on PWN-inoculated seedling LI 4. A, massive embolisms in the stem 1 cm above the inoculation wound. Arrows point to resin canals with crushed parenchyma cells in the embolized area. The arrowhead points to the annual ring border. B, patchy embolisms in cross section. The arrow points to a resin canal tylosoid. C, magnified view of tylosoid in a resin canal surrounded by water-filled tracheidslocated near the border of massive embolisms. The arrow points to an expanded epithelial cell. D, magnified view of an empty resin canal with destroyed epithelial cells. The arrowhead points to a nematode. Scale bars: A, 500 μm; B, 100 μm; C and D, 50 μm.

**Figure 6.**
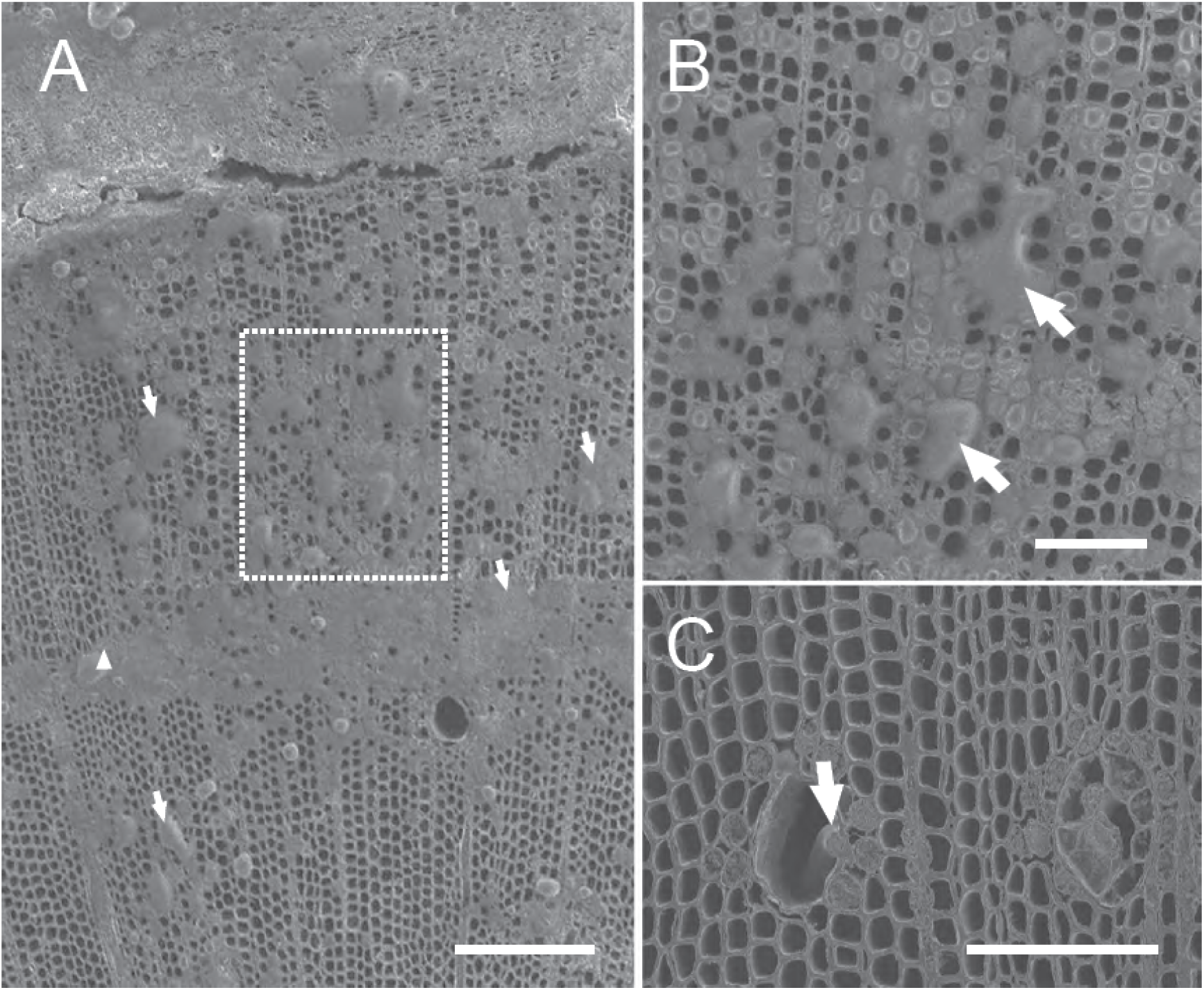
Cryo-SEM images of cross sections of 1-year-old stems 1 cm above the injection sites on PWN-inoculated seedling DI 1. A, mosaic-like distribution of resinous substances in xylem. Arrows point to the resinous substance in tracheids. The arrowhead points to the annual ring border. The dashed rectangle encloses the extended area in B. B, magnified view of the mosaic-like distribution of resinous substances in the xylem. Arrows point to resinous substances in the tracheids. C, magnified view of resin canals with destroyed epithelial cells. The arrow points to a nematode. Scale bars: A, 250 μm; B and C, 100 μm.

**Figure 7.**
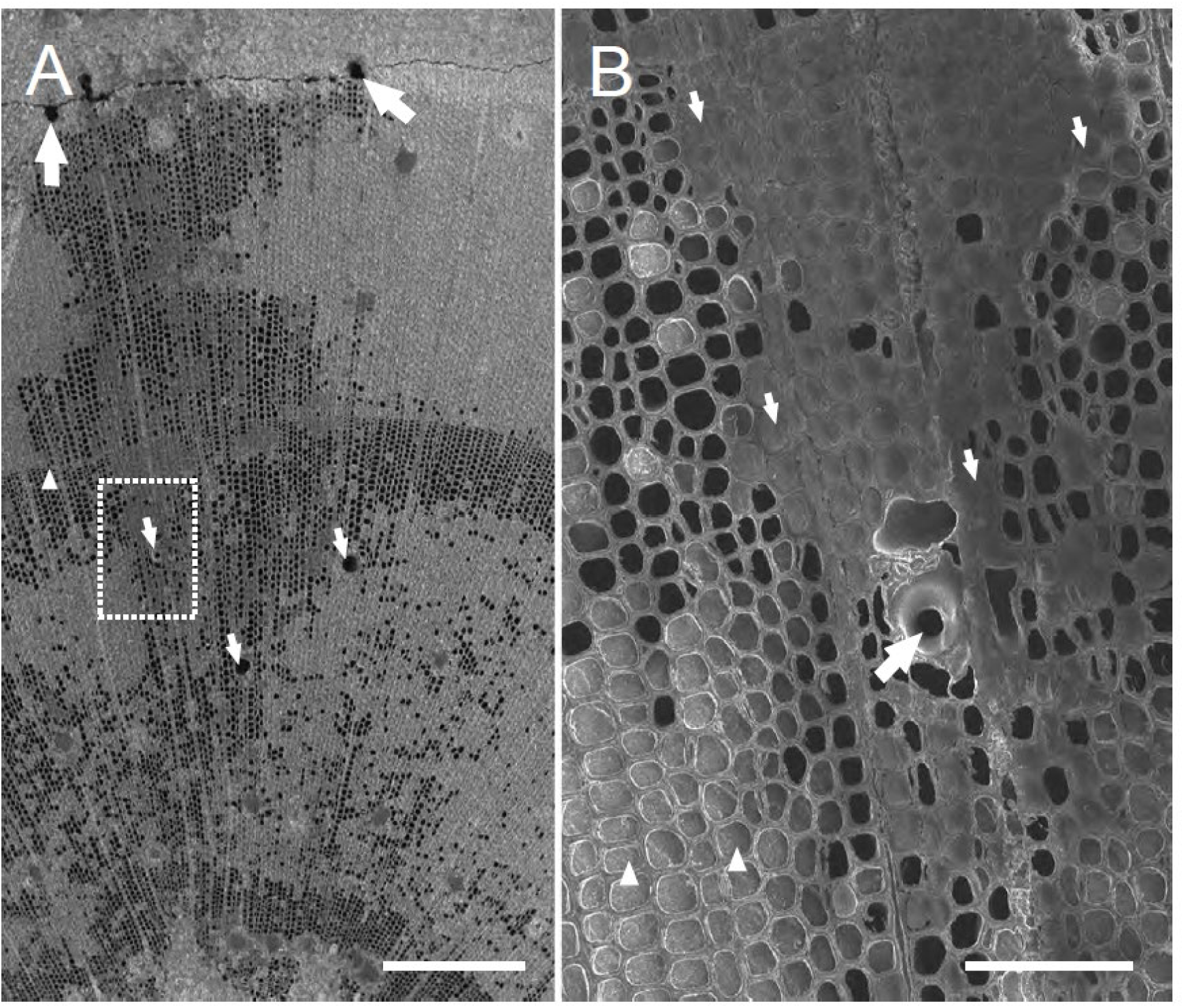
Cryo-SEM images of cross sections of 1-year-old stems 0.5 cm above the injection site on PWN-inoculated seedling DWI 1. A, massive embolisms 0.5 cm above the inoculation wound. The dashed rectangle encloses the extended area in B. Small arrows point to resin canals with destroyed epithelial cells. Large arrows point to cavities formed around the cambium. The arrowhead points to the annual ring border. B, magnified view of an axial resin canal in the 1-year-old xylem. The large arrow points to an axial resin canal partially filled with resin. Small arrows point to massive resinous substances. Arrowheads point to water in tracheids. Scale bars: A, 500 μm; B, 100 μm.

Most tracheids in PWN-infected seedlings that were not transpiring (DWI 3 and 4) were filled with water, but some near the inoculation wound or near the resin canals were plugged with resinous substances (Fig. 8A, B, D). We also observed tylosoid in the resin canals in several resin canals near the inoculation wound (Fig. 8A, C). Several juvenile PWNs and crushed living cells were observed in some resin canals containing resin (Fig. 8E).

**Figure 8.**
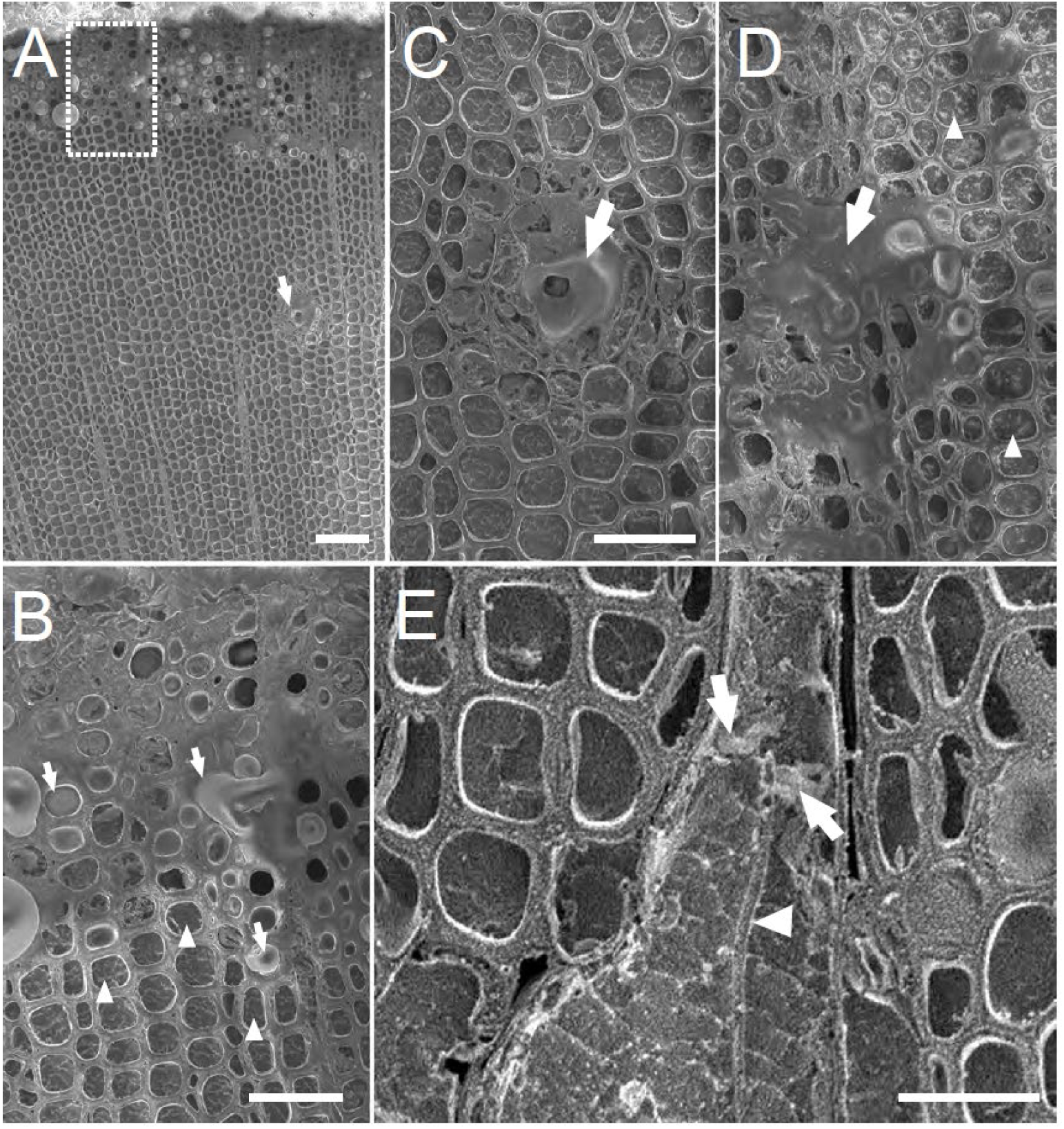
Cryo-SEM images of cross sections of 1-year-old stems 0.5 cm above the injection site on PWN-inoculated seedling DWI 3. A, water and resinous substance distribution 0.5 cm above the inoculation wound. The arrow points to a resin canal. The dashed rectangle encloses the extended area in B. B, magnified view of the inoculation site in current-year xylem. The arrowhead points to water in a tracheid. The arrows point to resinous substances in a tracheid. C, magnified view of tylosoid in a resin canal surrounded by water-filled tracheids. The arrow points to an expanded epithelial cell. D, transverse surface view of current-year xylem. The arrow points to a resin-filled tracheid. Arrowheads point to a water-filled tracheid. E, transverse surface view of a radial resin canal in current-year xylem. Arrows point to crushed living cells in the resin canal. The arrowhead points to a juvenile nematode. Scale bars: A, 100 μm; B–D, 50 μm; E, 25 μm.

## Discussion

We explored the clear differences between water stress-induced embolisms and abnormal embolisms in PWN-infected seedlings. How much pressure is required to induce abnormal embolisms at the inoculation wound in PWD? Japanese black pine seedlings generally have high water stress tolerance, and embolisms are rarely induced until −4.0 MPa has been reached (Umebayashi et al., 2016a). In contrast, abnormal patchy and massive embolisms were induced in PWN-infected seedlings under higher water potentials, as shown in previous studies (Fukuda, 1997; Kuroda, 1991; Yazaki et al., 2018). We first determined a negative threshold of −0.5 MPa for the occurrence of massive embolisms in specimens with PWD disease symptoms (Fig. 3B). Although we showed that massive embolization can be avoided by holding seedlings in darkness, thereby preventing leaf transpiration, the embolized area caused by a leakage of resin from resin canals into the surrounding tracheids gradually expanded in dark and/or wet conditions in nematode-infected trees (Figs. 1D, 6A, 7A, 8A). Thus, in PWN-infected trees, xylem embolisms can develop even in darkness without water stress, i.e., on rainy days. Many resin canals of the control trees were filled with resin (Fig. 4A). More importantly, we demonstrated that resin leaked from the resin canals to the tracheids decreased the surface tension, increasing the threshold water potential for cavitation up to −0.5 MPa, which is an unavoidably high level under field conditions. We showed that the occurrence of empty resin canals in infected trees is a result of resin leakage into tracheids.

In the developmental process of the embolized areas, radial expansion was related to the distribution of resin from the axial resin canals (Fig. 7). To invade inward from the inoculation wound, nematodes must migrate in the radial resin canals, and radially directed xylem embolisms are generally induced by PWN activity in the radial resin canals (Fig. 8E). Therefore, the developmental process of massive embolisms may be closely related to the distribution of radial and axial resin canals at the inoculation wound site. Kuroda et al. (1988) and Kuroda (1991, 2008) suggested that the occurrence of embolized tracheids may be related to the volatile exudate from ray parenchyma. However, we did not detect leakage from ray parenchyma to tracheids, and embolized tracheids were not distributed along the rays; they were distributed around resin canals and cavities. These observations can be explained by the occurrence of volatile exudates, including resin.

The detailed distribution of PWN around the inoculation wound has rarely been studied at the cellular level, although Kuroda (2008) believed that most nematodes in inoculation wounds are trapped in resin. PWNs were not observed in the resin canals with tylosoids in any of PWN-inoculated seedlings in our study (Figs. 5C, 8C); they were mostly found in the resin canals of crushed parenchyma cells (Figs. 4E, 5D, 6C, 8E). The resin canals of crushed parenchyma cells rarely occurred in patchy embolisms (Fig. 5B), but were frequent in massive embolisms (Fig. 5A, 6A, 7A). Thus, patchy embolisms can be induced by small numbers of migrating PWNs, but constant activity of many PWNs is required to induce the massive embolisms that can kill a pine tree by crushing parenchyma cells and inducing massive leakage of resin components into wide areas of the xylem. PWNs were usually observed in LI and DI seedlings, although several PWNs were also observed in DWI seedlings. Empty resin canals were detected in illuminated seedlings DWI 1 and 2, but not in seedlings DWI 3 and 4 held in darkness. The results are unclear, but PWN activity may depend on the condition of each host tree, and increases in the numbers of embolized tracheids may vary widely, as seen in LI and DWI seedlings.

The formation of tylosoid may not only contribute to the prevention of nematode migration around the inoculation wound, but may also facilitate the movement of resin in the resin canals. The crush of living cells by PWN may be generally induced at resin canal forming tylosoid as a result of the activities of PWN cellulase (Odani et al., 1985). After the destruction of living cells in the resin canals (Fig. 8E), the resin flows into the tracheids (in which the xylem sap is under negative pressure) where the living cells adjacent to tracheids around the resin canals are crushed. Once the resin in the resin canal has been sucked into the tracheids, the PWNs may readily move in and around the resin canals, and can also destroy the tylosoids. This may explain why tylosoids were rarely observed in massive embolisms and why nematodes have usually been observed in and around destroyed empty resin canals (Figs. 4E, 5D, 6C, Umebayashi et al., 2017). The succession of PWN activities and the pathological damage to the resin canals seem to be related to the development of massive embolisms. The quantity of leaked resin may depend on nematode activity, the formation of the tylosoids, and the distribution of crushed living cells; the distribution of leaked resin may be influenced by the xylem water potential. Strong negative pressure in daylight can cause cavitation and gas embolisms, but low negative pressure at night might also spread the distribution of leaked resin, leading finally to the occurrence of massive embolisms during the daytime. In future studies, we should examine the relationship between nematode distribution and anatomical changes in living cells at the inoculation wound site during early stages.

Although patchy embolisms were also observed around the inoculation site (Fig. 1C), leakage of resin from resin canals was barely detected in the embolized tracheids (Fig. 5B), as reported previously (Yazaki et al., 2018). Leakage was also barely detectable in the massive embolisms of the four LI samples (Figs. 4C, 5A). In these seedlings, negative pressure was stronger than in the DI and DWI seedlings, and embolisms with small amounts of leaked resin may have been present. The aspiration of bordered pits in these embolized tracheids may prevent further expansion of resin leakage to surrounding tracheids. This phenomenon should be investigated using three-dimensional analysis during the development of xylem dysfunction in PWD.

The development of the embolized area was slow in only one illuminated seedling (LI 4), in which patchy embolism abundances increased around the inoculation site (Fig. 1C). PWN-infected trees can survive if the embolized areas do not expand in the current-year xylem (Fukuda, 1997; Umebayashi et al., 2016b). The developmental speed and pattern of xylem embolisms may differ widely among samples, and further investigations should examine the detailed development of embolized areas in surviving seedlings.

## Conclusions

In summary, we examined the developmental process of xylem dysfunction during early stages of PWD by noninvasive CMRI and cryo-SEM. PWN activity induced tylosoid formation and promoted the crushing of living cells in/around resin canals. Xylem embolisms occurred at −0.5 MPa or lower, and the leakage of resin into tracheids occurred around resin canals (with the disappearance of tylosoid) and around the inoculation wound site. Thus, massive embolisms around the inoculation site were induced by a decrease in surface tension caused by the leaked resin. The massive embolisms expanded tangentially with threshold water potentials from −0.5 MPa to −0.7 MPa. Some of the leaked resin likely diffused axially during daylight hours, and embolized areas may develop axially. Extensive tangential diffusion may occur in darkness when the sap pressure gradient is relaxed, and the embolized zone may enlarge depending on the quantity of leaked resin.

## Funding information

The work was supported by KAKENHI (grant no. 23248022) from the Japan Society for the Promotion of Science (JSPS).

## Notes

### Competing Interest Statement

The authors have declared no competing interest.

