## Supplemental Fig. S1 for "Resin canal leakage underlying xylem dysfunction in pine wilt disease"

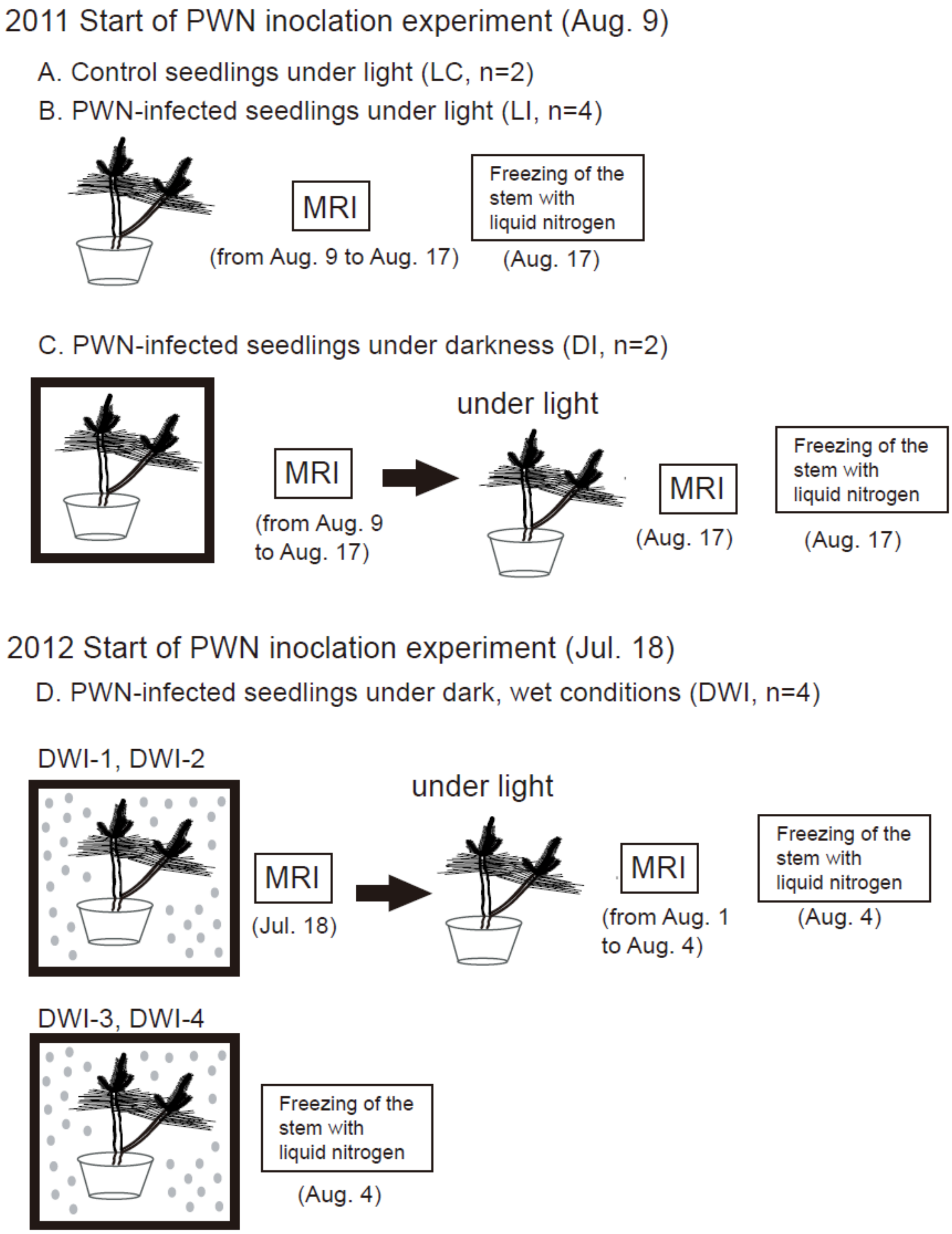


Figure S1. Schematic of the experimental design. Seedlings were held in our laboratory under light (LC and LI), in darkness by enclosing them in a blackout curtain to stop leaf transpiration (DI), or in darkness in a plastic bag with wet paper towel and misted with water (DWI).
